# Synaptic Zn^2+^ contributes to deleterious consequences of spreading depolarizations

**DOI:** 10.1101/2023.10.27.564408

**Authors:** Michael C. Bennett, Russell A. Morton, Andrew P. Carlson, C. William Shuttleworth

**Author notes:** Corresponding Author: C. W. Shuttleworth, Ph.D. Department of Neurosciences, MSC08 4740 1 University of New Mexico Albuquerque, NM 87131.

## Abstract

Spreading depolarizations (SDs) are profound waves of neuroglial depolarization that can propagate repetitively through injured brain. Recent clinical work has established SD as an important contributor to expansion of acute brain injuries and have begun to extend SD studies into other neurological disorders. A critical challenge is to determine how to selectively prevent deleterious consequences of SD. In the present study, we determined whether a wave of profound Zn^2+^ release is a key contributor to deleterious consequences of SD, and whether this can be targeted pharmacologically. Focal KCl microinjection was used to initiate SD in the CA1 region of the hippocampus in murine brain slices. An extracellular Zn^2+^ chelator with rapid kinetics (ZX-1) increased SD propagation rates and improved recovery of extracellular DC potential shifts. Under conditions of metabolic compromise, tissues showed sustained impairment of functional and structural recovery following a single SD. ZX-1 effectively improved recovery of synaptic potentials and intrinsic optical signals in these vulnerable conditions. Fluorescence imaging and genetic deletion of a presynaptic Zn^2+^ transporter confirmed synaptic release as the primary contributor to extracellular accumulation and deleterious consequences of Zn^2+^ during SD. These results demonstrate a role for synaptic Zn^2+^ release in deleterious consequences of SD and show that targeted extracellular chelation could be useful for disorders where repetitive SD enlarges infarcts in injured tissues.

## 1. INTRODUCTION

Spreading depolarizations (SDs) are waves of coordinated neuronal and glial depolarization that slowly propagate (2-4mm min^-1^) through neuronal tissue through mechanisms involving glutamate and/or K^+^ accumulation in the extracellular space (Leao, 1944; Somjen, 2001). The restoration of ionic gradients following SD is energetically demanding and, in metabolically challenged tissues, SD can be a primary mechanism of injury expansion (Hartings et al., 2017, 2003; von Bornstädt et al., 2015) Although there is much clinical interest in deleterious consequences of SD (Dreier et al., 2017; Hartings, 2017; Helbok et al., 2020; Lauritzen et al., 2011; Woitzik et al., 2013) there are currently limited treatments that target SD, with notable exceptions of agents such as ketamine and nimodipine (Carlson et al., 2020, 2018).

Synaptic release of glutamate release can be involved in the propagation of the SD waves (Mei et al., 2020; Van Harreveld, 1978). The divalent cation Zn^2+^ is co-stored and released together with glutamate from synaptic vesicles (Frederickson et al., 2005; Palmiter et al., 1996) and appears to regulate SD. Thus large waves of Zn^2+^ release and accumulation accompany SD (Carter et al., 2011), and the release of Zn^2+^ is implicated in slowing the progression of the propagating wave front, likely through inhibition of NMDA receptors (Aiba et al., 2012). We previously showed that experimental loading of very high levels of exogenous Zn^2+^ using an ionophore can impair recovery from SD (Carter et al., 2013). Persistent suppression of synaptic potentials after SD were due to accumulation of extracellular adenosine (Carter et al., 2013), consistent with the well-known inhibition of metabolic function by Zn^2+^ (Choi, 2020a; Dineley et al., 2003; Liu et al., 2021; Sensi et al., 2000). It is not yet known whether endogenous release of synaptic Zn^2+^ that occurs during the SD wave is sufficient to also contribute to deleterious consequences of SD.

Roles for synaptic Zn^2+^ release can be demonstrated experimentally by using mouse models with genetic deletion of the ZnT3 transporter (Anderson et al., 2015; Choi et al., 2020a; Cole et al., 2001). Pharmacological interventions that could be useful for clinical translation are complicated by multiple cellular sources of Zn^2+^ and a broad range of intracellular and extracellular actions. For example, large protein-bound intracellular Zn^2+^ pools regulate transcription and other essential functions (Palmiter et al., 1992). In the context of SD, an approach that chelates extracellular Zn^2+^ released from synapses during the propagating wave would be preferred over non-specific chelation in all intracellular and extracellular compartments. Capturing synaptically-released Zn^2+^ prior to intracellular accumulation is difficult with commonly-used chelators such as CaEDTA, due to their slow Zn^2+^ binding kinetics (Paoletti et al., 2009). To address this challenge, prior work has developed a Zn^2+^ chelator with rapid binding kinetics and has been used to demonstrate physiological synaptic regulation by Zn^2+^ (Pan et al., 2011).

The goal of the current study was to determine whether mobilization of endogenous Zn^2+^ can contribute to impaired recovery after SD. And if so, whether the source(s) of endogenous Zn^2+^ could be identified and targeted pharmacologically. In this study, we compare the effects of rapid extracellular Zn^2+^ chelation in control conditions and in a slice model of tissue vulnerability, where synaptic function is persistently suppressed after SD (Reinhart and Shuttleworth, 2018). We found that Zn^2+^ release from presynaptic terminals is sufficient to cause a substantial delay in recovery from SD, and this can be effectively prevented with a chelator with fast binding kinetics.

## 2. EXPERIMENTAL PROCEDURES

### 2.1. Animals and slice preparation

All animal procedures were performed in accordance with protocols approved by the UNM Health Sciences Center Institutional Animal Care and Use Committee. Adult (aged 6 – 8 weeks) male and female mice (both C57Bl/6 and/or ZnT3KO) were used for all experiments. Slice preparation was previously described (Shuttleworth et al., 2003). Briefly, animals were deeply anesthetized with a mixture of ketamine and xylazine (85 and 15 mg/mL, respectively), decapitated, and brains were removed into oxygenated ice-cold cutting solution (in mM): sucrose, 220; NaHCO3, 26; KCL, 3; NaH2PO4, 1.5; MgSO4, 6; glucose, 10; CaCl2, 0.2; equilibrated with 95%O2/5% CO2 and supplemented with 0.2mL ketamine (100mg/mL, Putney Inc., Portland, ME) to reduce excitotoxicity during the preparation of slices (Aitken et al., 1995). Brains were hemisected and cut in the coronal orientation (350μm slices) with a Leica VT1200 vibratome (Leica Biosystems, Deer Park, IL). Immediately after cutting, slices were incubated at 35°C for 60 min in artificial cerebrospinal fluid aCSF; containing (in mM): NaCl, 126; NaHCO3, 26; glucose, 10; KCl, 3; CaCl2, 2, NaH2PO4, 1.5; MgSO4, 1; equilibrated with 95% O2/5% CO2) then cooled to room temperature, for holding until transfer to the recording chamber.

### 2.2. SD generation

Slices were individually transferred into a submersion recording chamber with nylon slice supports (RC-27L, Warner Instruments, Holliston, MA) either to allow for aCSF flow on both sides of the slice or positioned to restrict flow to only one side of the slice. The restricted configuration reduces metabolic capacity and models vulnerable conditions as previously described (Reinhart and Shuttleworth, 2018). Slices were continuously superfused with oxygenated aCSF (95% O2/95% CO2) at ∼2ml/min. Bath temperature was maintained at 32°C. To evoke SDs, pressure microinjection (50 ms, 30 psi; UniBlitz Shutter D122, Artisan Technology Group, IL, USA) of KCL (1M) was used via glass micropipette placed 50μm below the surface in the stratum radiatum of CA1 region of hippocampal tissue. Except where specified, single SDs were initiated in each slice. Intrinsic optical signals (IOS) of brain slices using trans-illumination of visible light (≥ 600 nm) was used to monitor SD initiation and propagation as well as tissue recovery as described previously (Obeidat and Andrew, 1998; Reinhart and Shuttleworth, 2018) via 4X objective (Olympus, 0.10 NA). IOS data was captured using a cooled CCD camera (Imago, Till Photonics) and then analyzed using TillVision software (TillPhotonics, version 4.01). Transmitted light was normalized to baseline and IOS was expressed as a percent change in transmission of visible light (ΔT/T0 × 100).

### 2.3. Electrophysiology

Excitatory postsynaptic potentials (EPSPs) were generated by stimulation of Schaffer collaterals in stratum radiatum of area CA1 with concentric bipolar stimulating electrodes (FHC, Bowdoin, ME, USA) and recordings were measured using microelectrodes placed 250μm from the stimulating electrode and 50μm below tissue surface. Test pulses (50μs, 0.1Hz, 0.5-2.5mA) were set at 50 – 75% maximum EPSP amplitude. SD duration was measured from peak maximum to 80% recovery of the extracellular DC potential shift. Signals were analyzed using Clampfit 11.1 software (Molecular Devices, Sunnyvale, CA, USA).

### 2.4. Fluorescence imaging

Extracellular Zn^2+^ transients were assessed using a fluorescent indicator (FluoZin-3) as previously described (Carter et al., 2011). Briefly, FluoZin-3 (Na^+^ salt) was dissolved in ACSF (2μM) and superfused over slices (C57Bl/6 or ZnT3KO) at 32°C. CaEDTA (1mM) was included to reduce background fluorescence and allow for analysis of the rapid phase of Zn^2+^ accumulation (Carter et al., 2011). FluoZin-3 was excited at 495nm and emission detection at 535/50nm recorded by using a 2 Hz monochromator/CCD imaging system (Polychrome V, Till Photonics GmbH, NY; Imago, Till Photonics, Rochester, NY, USA).

### 2.5. Reagents

Both TPEN (100mg powder) and CaEDTA (100g powder) were purchased from Sigma-Aldrich (St. Louis, MO, USA), and dissolved in DMSO (25 mM stock) or ACSF (1mM), respectively. ZX-1 (100 mg powder) was purchased from Strem (Newburyport, MA, USA) and dissolved in water (100mM stock). CaEDTA was made up daily, while ZX-1 and TPEN were diluted to experimental concentration daily (100μM and 50μM, respectively) from stock.

### 2.6. Statistical analysis

Data are reported as mean ± SEM. Statistical analyses (repeated measure one-way analysis of variance (ANOVA), paired and unpaired t-tests) were calculated using GraphPad Prism (version 10.00; La Jolla, CA). Statistical significance was determined by P values < 0.05, with Bonferroni correction during multiple comparisons.

## 3. RESULTS

### 3.1 Rapid Zn^2+^ chelation can alter SD characteristics in healthy tissues

ZX-1 is a Zn^2+^ chelator with kinetics rapid enough to inhibit actions of Zn^2+^ released at synapses (Anderson et al., 2015; Pan et al., 2011). Figure 1 shows the effects of ZX-1 on characteristics of SD generated by focal microinjections of K+, in nominally healthy conditions (see Methods). Under these recording conditions, SDs can be generated repetitively at 15 min intervals, with full recovery between events. SD propagation was monitored from intrinsic optical signals (Fig 1A) and extracellular potential shifts corresponding to depolarization and recovery during SD (“DC shifts”) were recorded with a microelectrode placed in stratum radiatum (Fig 1B). ZX-1 significantly increased SD propagation rate (Fig 1C) and also decreased the duration of DC shifts (Fig 1C). Both effects were partially reversed following ZX-1 washout.

**Figure 1:**
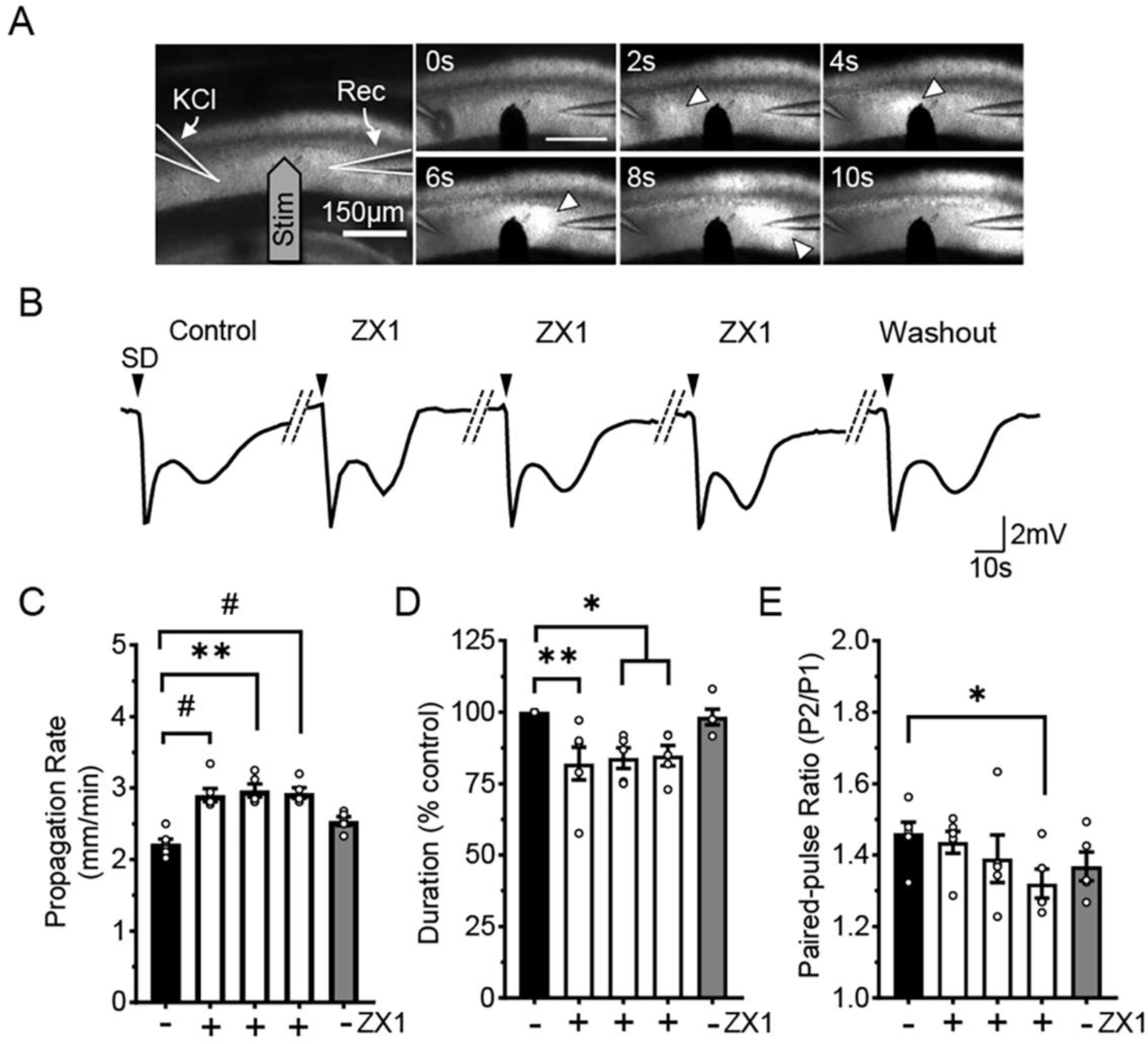
ZX-1 accelerates SD propagation and recovery. **A:** Initiation and propagation of a single SD event. Left most panel shows the location of recording electrode in a transmitted light image. SD was initiated by focal KCl microinjection (*KCl*). Visualization of SD was obtained through imaging the slowly propagating wave of increased light transmission (white arrowheads). A second microelectrode (*Rec*) was used to record DC potential shifts characteristic of SD, and also field EPSPs following Schaffer collateral stimulation via bipolar electrode (*Stim*). **B:** DC shift recordings of SD, during a sequence of 5 events generated at 15 min intervals in the same slice, prior to (*first trace*) during ZX-1 (*three middle*) and after ZX-1 washout (*last trace*). *Black arrowheads* indicate SD initiation. **C-E:** Summary data from experiments shown in A and B (n=5), demonstrating increased propagation rates (**C**), decreased duration of DC shift (**D**), and also decreased paired-pulse ratio from pairs of EPSPs generated with 50ms inter-pulse intervals (**E**). *\*P* < 0.05, *\*\*P* < 0.005, *^#^P* < 0.0005.

One explanation for reduced duration of DC shifts could be improved metabolic capacity of tissues (Lindquist and Shuttleworth, 2014; Reinhart and Shuttleworth, 2018), caused by reduction of Zn^2+^ loading during SD. Figure 1E provides initial support for this hypothesis, by showing that paired-pulse ratio (PPR) of excitatory postsynaptic potentials (EPSPs) is reduced during ZX-1 exposures. PPR was reduced prior to the onset of SDs, consistent with decreased cumulative adenosine-dependent suppression of presynaptic release probability (Lindquist and Shuttleworth, 2017, 2012) during the series of SDs in ZX-1. Figure 2 shows that ZX-1 also significantly increased recovery of EPSPs following a single SD. As previously described, EPSPs are almost abolished after SD and recovery relatively slowly over the next 5-10min as a consequence of metabolic depletion and activation of presynaptic adenosine A1 receptors (Lindquist and Shuttleworth, 2014, 2012). The increased recovery rate of EPSPs in ZX-1 is therefore consistent with decreased metabolic demand in the recovery phase after each SD.

**Figure 2:**
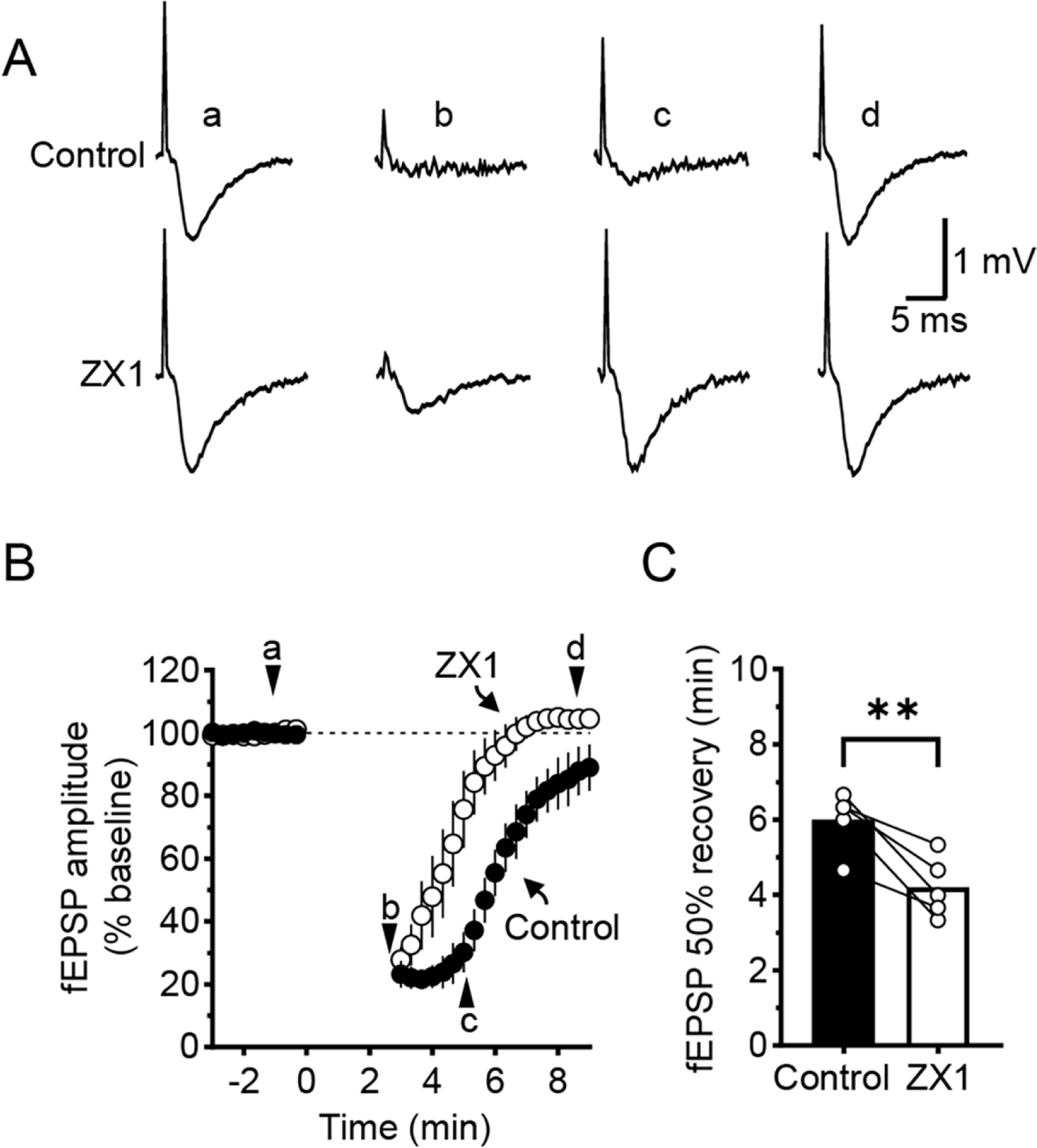
ZX-1 accelerates recovery of excitatory postsynaptic potentials (fEPSPs). **A:** Effect of SD on representative fEPSPs generated in control conditions (*top*) and during ZX-1 exposure (*bottom*). Timepoints a-d refer to times indicated below in panel B, before, during, and after recovery from SD. **B:** Time course of fEPSP suppression and recovery following SD in control (*black circles*) and ZX-1 (*white circles*). **C:** Summary data showing time to 50% recovery of fEPSP amplitude after SD (n = 5), *\*\*P* < 0.01.

### 3.2 ZX-1 improves recovery from SD in metabolically compromised tissues

SDs are fully recoverable in tissues with adequate metabolic substrate availability, but can lead to irreversible neuronal failure under conditions of metabolic compromise (Reinhart and Shuttleworth, 2018). We therefore next tested whether rapid chelation of Zn^2+^ could be sufficient to protect neurons from persistent impairment from SD in slices that were metabolically restricted. A decrease in metabolic supply was achieved using flow restriction as previously described (see Methods) and SDs initiated and recorded as above in Figure 1. Under these conditions, vulnerability to SD is readily observed in IOS recordings showing persistent lack of recovery following SD. Figure 3A compares normally recovering IOS signals in control conditions with the persistent decrease in light transmission observed at late time points after SD. Prior work has associated IOS decreases after SD with tissue compromise (Obeidat and Andrew, 1998; Reinhart et al., 2021; Reinhart and Shuttleworth, 2018). ZX-1 prevented the negative-going IOS signal after SD in vulnerable conditions, and significantly improved signals at 10 min post SD. Figure 3C shows that non-selective intracellular and extracellular Zn^2+^ chelation with TPEN (50μM) was similarly effective, but that an extracellular chelator with slower kinetics (CaEDTA, 100μM) was unable to prevent persistent IOS decreases.

**Figure 3:**
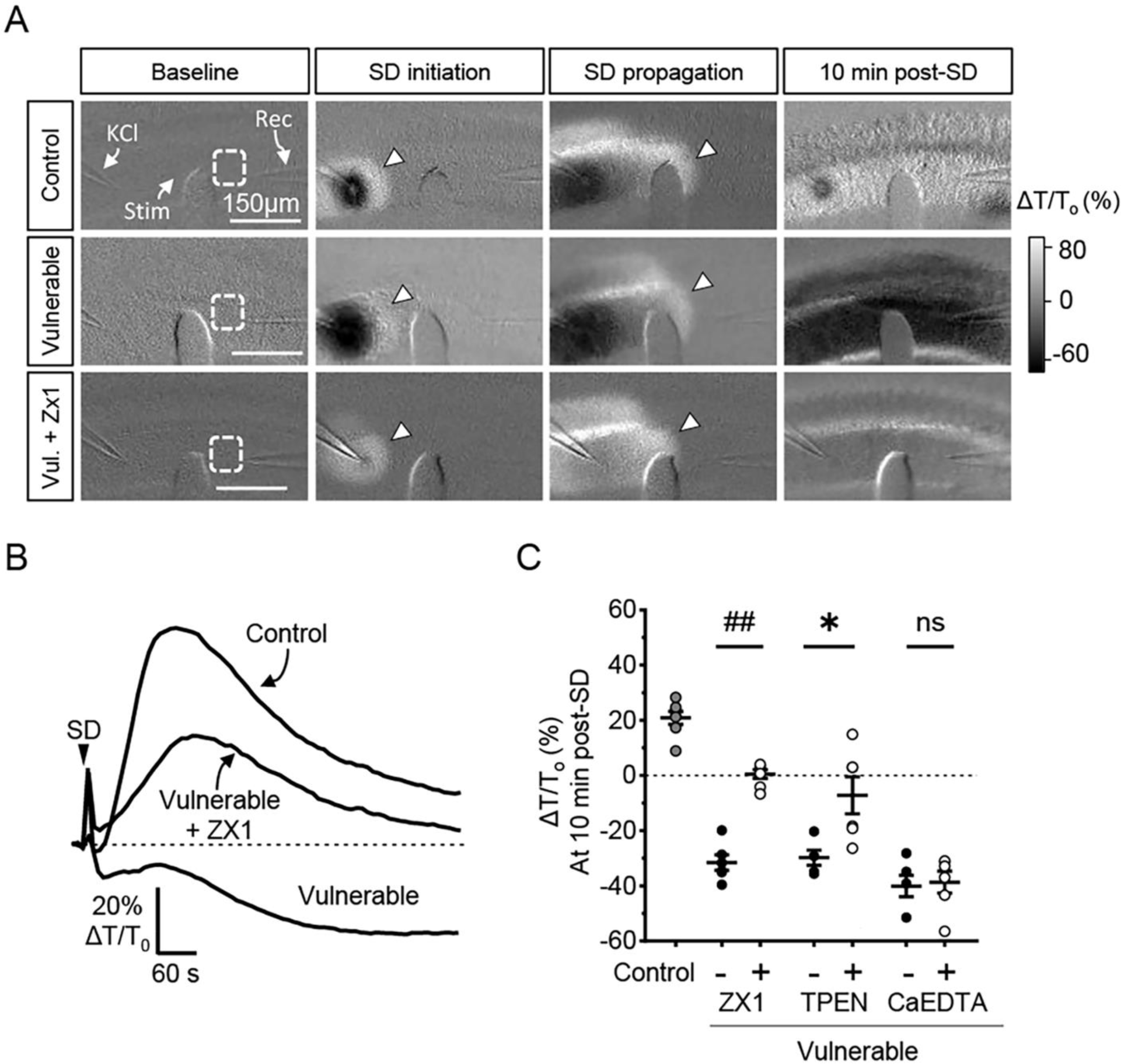
ZX-1 improves recovery from SD in metabolically compromised slices. **A:** Representative intrinsic optical signals (IOS) showing SD propagation and recovery. Responses in control recording conditions (top) are compared with metabolically compromised recording conditions (middle, *Vulnerable,* see Methods). The bottom panels show that ZX-1 markedly improved recovery of IOS after SD in the vulnerable conditions (bottom row). **B:** IOS signals from the representative recordings in A, with data extracted from the white dotted region (*ROI*) in baseline images. **C:** Summary data of effects of three different Zn^2+^ chelators (ZX-1, TPEN, CaEDTA) on IOS recovery after SD in vulnerable conditions. Recovery in control conditions is also shown for reference. \**P* = 0.0172, ^##^*P* < 0.0001

Consistent with improved structural recovery, as indicated by IOS signals, ZX-1 also improved functional recovery of EPSPs. Figure 4 compares EPSP recovery in the three conditions (control, vulnerable and vulnerable + ZX-1). As previously shown, FEPSP recovery was greatly prolonged in vulnerable conditions. ZX-1 pre-exposure resulted in significantly improved rate of recovery, with EPSP amplitude being significantly increased at 20min post SD. Together, these findings suggest that endogenously-mobilized Zn^2+^ contributes to the metabolic burden to tissues recovering from SD, and that rapid chelation can be sufficient to prevent persistent impairment in vulnerable tissues.

**Figure 4:**
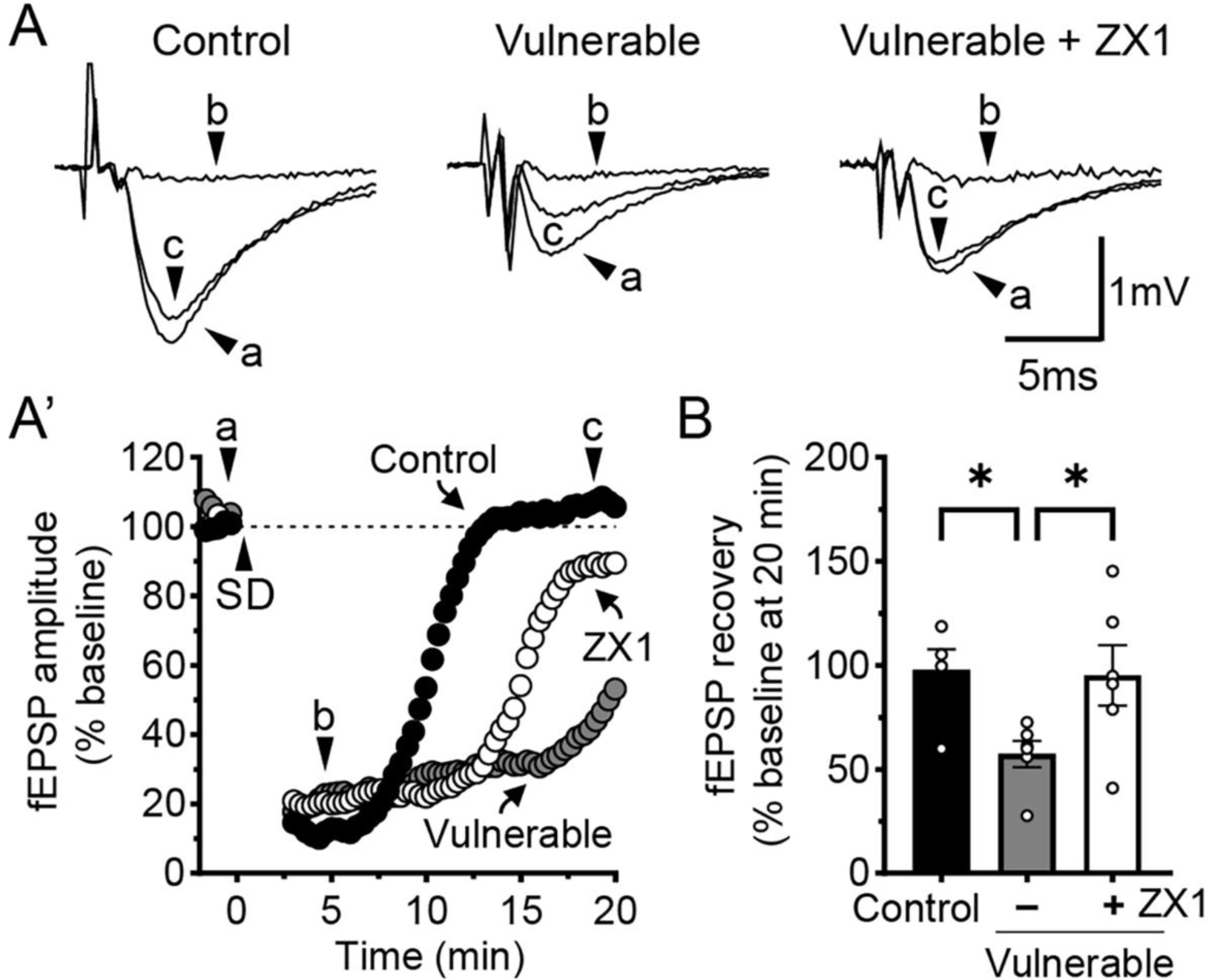
ZX-1 improves synaptic recovery in metabolically compromised slices. **A:** Effect of SD on representative fEPSPs generated in either control conditions (left) or in the metabolically compromised model (middle and right). Time points (a-c) refer to times illustrated in plot below (A’). fEPSP recovery was impaired in the vulnerable conditions, and significantly improved by ZX-1 in these conditions. **B:** Summary data showing EPSP recovery 20 minutes after SD initiation. ZX-1 significantly improved recovery compared to un-treated vulnerable slices (*grey bar*). *\*P* < 0.05

### 3.3 Synaptic Zn^2+^ release during SD can be attenuated by ZX-1 chelation and ZnT3 knockout

We next examined whether release of Zn^2+^ from presynaptic vesicles is a likely source of detrimental Zn^2+^ effects in vulnerable tissues. Initial experiments tested whether ZX-1 was able to chelate synaptic Zn^2+^ released during the advancing SD wavefront. Figure 5 shows measures of extracellular Zn^2+^ accumulation, detected with the fluorescent indicator FluoZin3. In vulnerable tissues, SD generated propagating waves of extracellular Zn^2+^ accumulation, similar to reported previously with healthy brain slices (Carter et al., 2011). Zn^2+^ increases were observed prominently at the site of focal K^+^ application, before propagating with the advancing SD wavefront. Figure 5B shows that ZX-1 effectively reduced the SD-associated Zn^2+^ increases. Extracellular increases at the SD initiation site was partially reduced (by ∼30%), but signals at the propagating wavefront were more effectively reduced (by ∼75%). Under the same recording conditions of metabolic compromise, genetic ablation of synaptic Zn^2+^ release abolished Zn^2+^ signals, as previously reported in healthy tissues (Carter et al., 2011). Taken together with effectiveness of ZX-1, these findings suggest that synaptic release could be the source of Zn^2+^ that leads to persistent neuronal compromise in compromised tissues.

**Figure 5:**
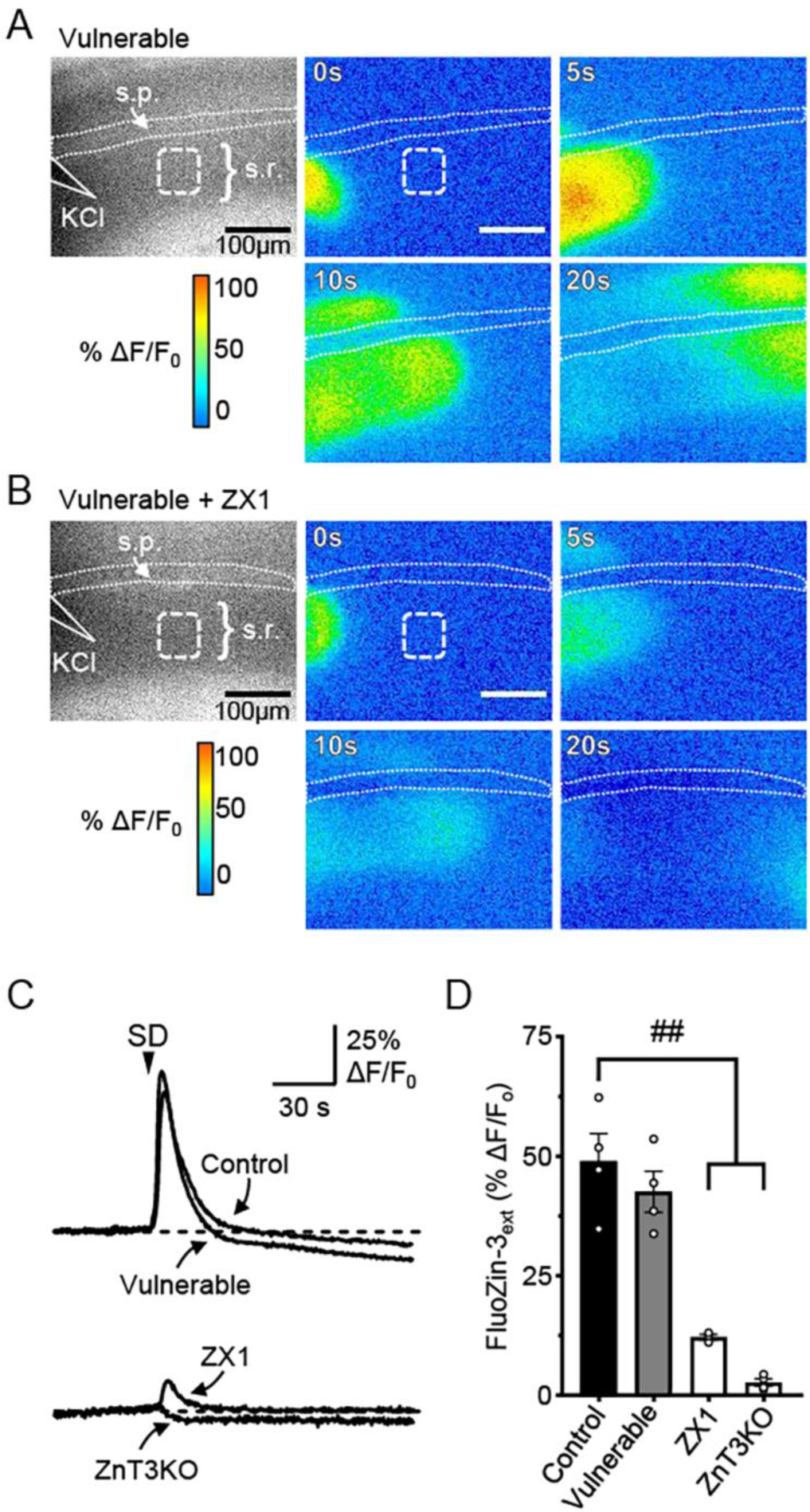
Synaptic release of Zn2+ and chelation with ZX-1. **A:** Extracellular Zn^2+^ accumulation detected with FluoZin-3 during SD initiation and propagation in vulnerable conditions. The first panel shows the location of focal *KCl stimulus* and subsequent pseudocolor panels show FluoZin-3 increases at times indicated following SD initiation. Zn^2+^ increases are prominent at both the initiation site (0-5s) and tracking with the spreading of the SD wave (10-20s). **B:** ZX-1 greatly decreased FluoZin-3 signals associated with SD propagation. **C:** Kinetics of FluoZin-3 signals from the propagating wavefront of SD under different recording conditions. Top panel compares transients obtained in control and vulnerable conditions, showing very similar amplitudes and kinetics. The bottom panel shows that ZX-1 greatly reduces evoked transients, and that genetic ablation of presynaptic Zn^2+^ transporter ZnT3KO abolishes SD-evoked increases. **D:** Summary data showing the significant decrease in SD-evoked FluoZin-3 transients in both ZX-1 (n=3) and ZnT3KO (n=4) compared to control (n=4) and vulnerable (n=4) controls. ^##^*P* < 0.0001

Consistent with these results, Figure 6 shows that genetic removal of synaptic Zn^2+^ is sufficient to improve structural and functional recovery of tissues after SD. Figure 6 studies were all completed with metabolically compromised conditions, and show that recovery of fEPSPs after SD was comparable to the beneficial effects observed above with ZX-1 (Figure 6A,D and compare with Figure 4). Structural recovery was also substantially improved, as indicated by lack of sustained decrease in IOS signals (Figure 6C). Finally, it was noted that SD propagation rate was also enhanced in the vulnerable conditions, consistent with a role for synaptic Zn^2+^ in limiting SD propagation in these conditions of limited substrate availability.

**Figure 6:**
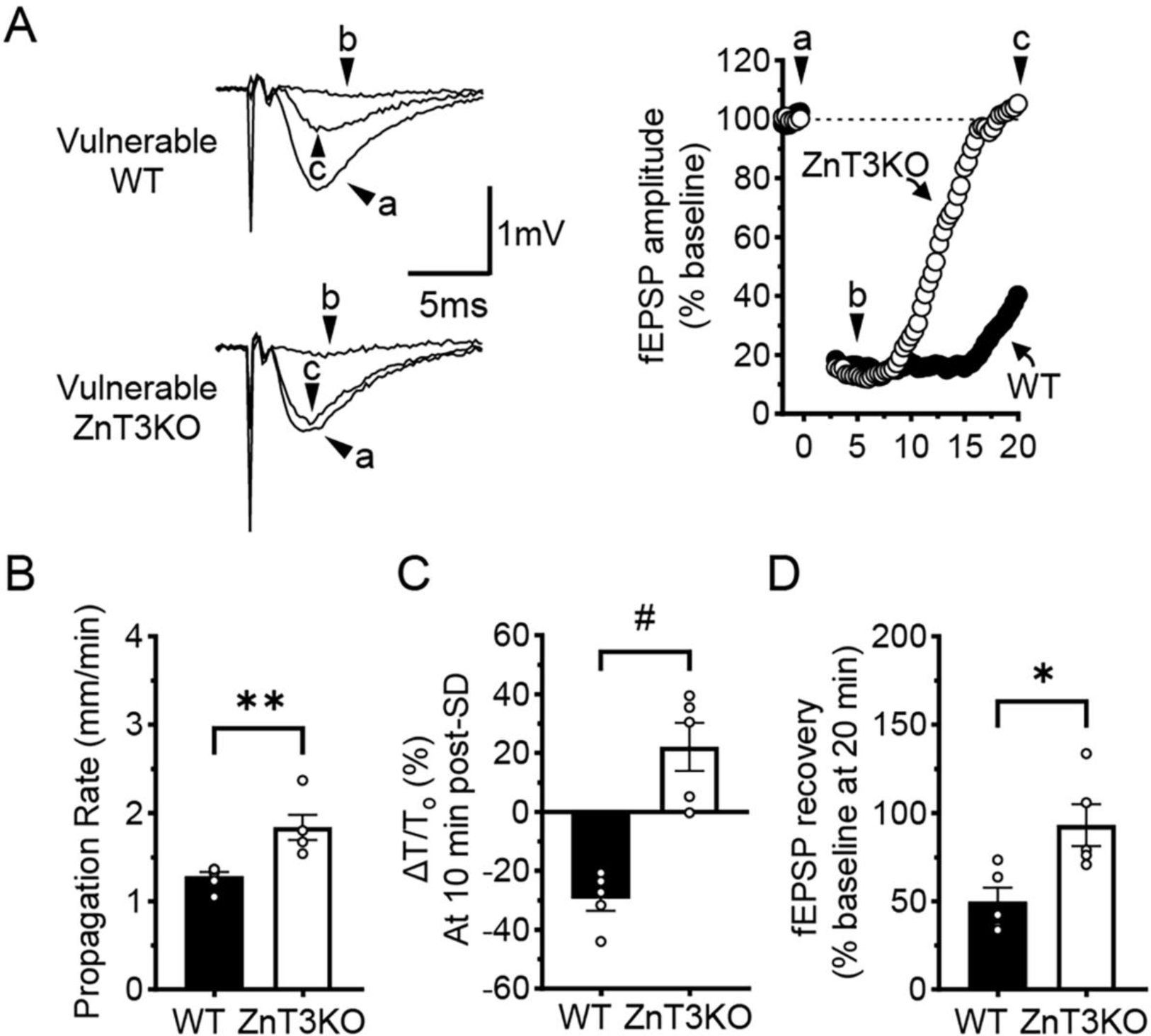
ZnT3KO improves recovery after SD in metabolically compromised tissue. **A:** Effect of SD on representative fEPSPs generated in either wild type controls (top) or in ZnT3KO slices (bottom) in the metabolically compromised model. Time points (a-c) refer to times illustrated in the right plot. **B-D:** Summary data of effects of ZnT3KO on propagation rate of SD through the imaging field (**B**), recovery of IOS transmittance (**C**), and recovery of fEPSP amplitudes (**D**) after SD in vulnerable model. Wild type controls in vulnerable are also presented for reference. \**P* < 0.02, \*\**P* < 0.003, ^##^*P* < 0.0005

## 4. DISCUSSION

### 4.1. General

The main findings of the present study are 1) the wave of synaptic Zn^2+^ release that occurs during SD can contribute to deleterious consequences of SD in vulnerable brain tissue and 2) these effects can be prevented by extracellular chelation with a rapidly-acting chelator. These observations suggest an opportunity to pharmacologically target Zn^2+^ an adjunct approach to limit injury following SD in a range of acute brain injuries.

### 4.2. Synaptic release of Zn^2+^ during SD

It appears that Zn^2+^ is co-released with glutamate from synaptic vesicles during the propagation of SD. Extracellular accumulation of Zn^2+^ is detected as a wave that matches intrinsic optical signal propagation of SD through brain slice preparations (Carter et al., 2011) and fluorescence detection of extracellular glutamate accumulation with iGluSnFr shows a similar or identical pattern (Enger et al., 2015). Deletion of ZnT3 abolishes zinc uptake into glutamatergic vesicles in excitatory neurons (Palmiter et al., 1996) and blocks extracellular Zn^2+^ waves during SD. SD-induced Zn^2+^ waves were characterized previously with focal K^+^ stimulation in healthy tissues (Carter et al., 2011) and in the present studies with partial metabolic compromise the patterns of accumulation appeared very similar. The fact that ZX-1 greatly reduced SD-evoked extracellular Zn^2+^ waves shows that this chelator is able to sequester synaptically released Zn^2+^ under these conditions (Figure 5). It is possible that the kinetics of uptake after SD may be modified by decreased energy availability, but were unable to be detected because of the use of a slow chelator (CaEDTA) in fluorescence imaging studies with FluoZin-3 to increase detection sensitivity (Carter et al., 2011; Qian and Noebels, 2005).

While there has been a focus on the hippocampus for synaptic Zn^2+^ biology, Zn^2+^ is also found in neocortical circuits in significant levels (Frederickson et al., 2005) and release is seen during SD propagations in murine neocortex in vivo (R.E. Carter and C.W. Shuttleworth, unpublished observations). It is thus likely that similar Zn^2+^ accumulation follows SD waves that occur in the neocortex and contribute to neuronal injury following stroke and trauma. Whether or not SD-evoked Zn^2+^ waves occur in the human neocortex has not been directly tested, but this would appear likely based on the presence of synaptic Zn^2+^ in these tissues (Frederickson et al., 2005) and the conservation of cellular mechanisms of SD between species (Somjen, 2001).

### 4.3 Damaging effects of Zn^2+^ accumulation

An extensive literature demonstrates that Zn^2+^ accumulation can lead to neuronal injury or death (Choi et al., 2020a; Frederickson et al., 2005; Liu et al., 2021). Intracellular actions of Zn^2+^ on metabolic function are often considered primary sites of action for deleterious effects. Zn^2+^ can bind to GAPDH to inhibit glycolysis and can also lead to activation of PARP-1 resulting in downstream DNA damage and consumption of NAD^+^ (Choi et al., 2020a; Kim and Koh, 2002; Sheline et al., 2000). Zn^2+^ loading of disruption of mitochondrial function has also been strongly linked to neuronal injury (Dineley et al., 2003; Liu et al., 2021; Medvedeva et al., 2022, 2009; Sensi et al., 2000). In the present study, Zn^2+^ chelation with ZX-1 improved recovery of DC shifts that occur during SD, which is one indicator of improved metabolic capacity by removal of the Zn^2+^ wave (Figure 1). Likewise, recovery of evoked synaptic potentials was improved by ZX-1 in healthy tissues, consistent with reduced metabolic depletion and adenosine accumulation during the ∼10 min following SD (Figure 2). The effects of Zn^2+^ removal were more profound in the vulnerable tissues, as would be expected if Zn^2+^ accumulation impaired neuronal metabolism. Thus when oxygen and glucose availability was limited in the vulnerable model, SD alone caused a long-lasting suppression of synaptic potentials (Figure 4) and a characteristic persistent decrease in intrinsic optical signals that have been associated with neuronal injury (Obeidat and Andrew, 1998). Preventing Zn^2+^ waves with either genetic deletion of ZnT3 or with ZX-1 chelation both produced substantial recovery of both structural and functional deficits. These results imply a significant role for synaptic Zn^2+^ accumulation in the cascade of damaging events that occur following SD in models of tissue vulnerability.

Multiple ion channels and transporters expressed on neurons are capable of increasing postsynaptic Zn^2+^ accumulation following SD. For example, Zn^2+^ flux via L-type calcium channels and NMDA receptors can be significant (Kerchner et al., 2000; Marin et al., 2000), and both these channel types contribute to ionic loading after SD (Aiba and Shuttleworth, 2012; Dietz et al., 2008). Intracellular Ca^2+^ loading is well established to contribute to neuronal injury following excessive glutamate stimulation (Choi, 2020b), and is implicated in neuronal damage following SD in compromised tissues (Aiba and Shuttleworth, 2012). Zn^2+^ and Ca^2+^ can both contribute to the initiation of SD and actions of Zn^2+^ can serve as an upstream regulator of excitotoxic Ca^2+^ loading (Dietz et al., 2009; Shuttleworth and Weiss, 2011; Vander Jagt et al., 2009). For these reasons, targeting Zn^2+^ accumulation could be a complementary strategy for limiting Ca^2+^-dependent neuronal injury following episodic glutamate surges during SD (Hinzman et al., 2015).

Zn^2+^ accumulation via a range of Zn^2+^ transporters are also expressed on neurons and could contribute to Zn^2+^ accumulation after SD. Flux through these different pathways has generally been characterized under resting conditions or following synaptic stimulation, and little is known of relative contributions during the extreme stimulation conditions of SD. During SD, neurons can undergo sustained depolarization for a minute or more, with severe disruptions of ionic gradients and energy depletion. How these factors modify Zn^2+^ transport in the wake of SD is currently unknown. In addition to neurons, Zn^2+^ could deplete metabolism in astrocytes (Lee and Koh, 2010), which also contribute to regulation of SD propagation and recovery (Seidel et al., 2016). Understanding these sources and sites of Zn^2+^ action could be helpful in refining Zn^2+^ targeting strategies in conditions where SD contributes to injury progression.

### 4.4 Beneficial effects of Zn^2+^

Synaptic Zn^2+^ release acts extracellularly to limit the rate of SD propagation through neuronal tissue (Aiba et al., 2012) most likely due to its inhibitory effect on NMDA receptors containing the GluN2A subunit (Anderson et al., 2015; Paoletti et al., 1997). It has been suggested previously that extracellular Zn^2+^ application may be beneficial to tissue as it can inhibit SD propagation (Aiba et al., 2012). Interestingly, Zn^2+^ inhibitory effects are dependent on tissue metabolic state as this sensitivity to Zn^2+^ can be completely lost in SDs initiated in hypoxic tissue (Aiba and Shuttleworth, 2013). In the current study, synaptic Zn^2+^ contributed to slowing the rate of propagation metabolically compromised tissues (Figure 6B), suggesting that under these conditions of partial hypoxia/hypoglycemia, the inhibitory effects of Zn^2+^ on NMDA or other redox-modulated targets is still maintained. This suggests that in ischemic penumbra, beneficial effects of Zn^2+^ on SD propagation could be maintained. However, it appears that these effects can be far outweighed by deleterious consequences of Zn^2+^ on increasing the metabolic burden of SDs and persistent lack of structural and functional recovery after SD in vulnerable tissues.

### 4.5 Potential translation

As described above (Introduction), inhibitors of NMDA receptors and L-type Ca^2+^ channels (ketamine, memantine, nimodipine) are being tested as therapeutic agents to target SDs. These agents, used for normal clinical care (sedation in the case of ketamine and vasospasm prevention in aneurysmal subarachnoid hemorrhage in the case of nimodipine), have been demonstrated to decrease SD in pre-clinical models (Dreier et al., 1998; Reinhart et al., 2023; Reinhart and Shuttleworth, 2018; Sánchez-Porras et al., 2022; Tóth et al., 2020) and also suggested in clinical studies (Carlson et al., 2020, 2018; Hertle et al., 2012). While inhibition of Ca^2+^ accumulation is generally considered to underly beneficial effects of these agents, the discussion above implies that inhibition of Zn^2+^ accumulation after SD could also contribute. The relative contributions of Zn^2+^ and Ca^2+^ could theoretically be determined in future studies by comparison with selective removal of Zn^2+^ after SD. There have been prior clinical studies of Zn^2+^ chelation, most notably with the membrane activated chelator DP-b99. This agent demonstrated promise in terms of stroke neuroprotection by modulating calcium, zinc and other ions within cell membranes (Diener et al., 2008). A phase 3 clinical trial however failed to show a benefit in clinical outcomes (Lees et al., 2013). This and other failed stroke neuroprotection studies may be limited by a lack of prior understanding of the role of SD in stroke and other brain injuries. SD leads to discrete episodes of glutamate and Zn^2+^ release and excessive accumulation that occur intermittently over a period of hours and days following insult, in contrast to prior assumptions of continuous release from an infarct core (Hartings et al., 2017). Strategies to target the patients at most risk of secondary expansion related to SD, or to specifically target SD or its consequences may therefore re-open windows into previously tested neuroprotective strategies based on a more complete understanding of the physiology of stroke expansion.

## ACKNOWLEDGEMENTS

Supported by NIH grants NS129351, NS106901 & GM109089. Author contributions: MCB designed and performed experiments, analyzed data and co-wrote manuscript, RAM contributed to design of experiments and edited manuscript, APC contributed to design of experiments, interpretation and editing of manuscript, CWS designed experiments, interpreted data and co-wrote manuscript.

